# Thinking outside the CaaX-box: an unusual reversible prenylation on ALDH9A1

**DOI:** 10.1101/2022.08.29.505770

**Authors:** Kiall F. Suazo, Garrett L. Schey, Shelby A. Auger, Ling Li, Mark Distefano

## Abstract

Protein lipidation is a post-translational modification that confers hydrophobicity on protein substrates to control their cellular localization, mediate protein trafficking, and regulate protein function. In particular, protein prenylation is a C-terminal modification on proteins bearing canonical prenylation motifs catalyzed by prenyltransferases. Such types of proteins have been of interest owing to their potential association with various diseases. Chemical proteomic approaches have been pursued over the last decade to define prenylated proteomes (prenylome) and probe their responses to perturbations in various cellular systems. Here, we describe the discovery of prenylation of a non-canonical prenylated protein, ALDH9A1, which lacks any apparent prenylation motif. This enzyme was initially identified through chemical proteomic profiling of prenylomes in various cell lines. Metabolic labeling with an isoprenoid probe using overexpressed ALDH9A1 reveals that this enzyme can be prenylated inside cells but does not respond to inhibition by prenyltransferase inhibitors. Site-directed mutagenesis of the key residues involved in ALDH9A1 activity indicate that the catalytic C288 bears the isoprenoid modification likely through an NAD^+^-dependent mechanism. Furthermore, the isoprenoid modification is also susceptible to hydrolysis, indicating a reversible modification. We hypothesize that this modification originates from endogenous farnesal or geranygeranial, the established degradation products of prenylated proteins and results in a thioester form that accumulates. This novel reversible prenoyl modification on ALDH9A1 expands the current paradigm on protein prenylation by illustrating a potentially new type of protein-lipid modification that may also serve as a novel mechanism for controlling enzyme function.

## 1. Introduction

Protein prenylation is an essential post-translational modification (PTM) required for certain proteins to properly localize in cellular membranes and for mediating protein-protein interactions and protein trafficking.^1^ In this type of PTM, a farnesyl or geranylgeranyl group(s) is irreversibly appended through a thioether linkage on a cysteine residue near the C-terminus of a protein. There are currently four characterized prenyltransferases that catalyze such modifications. Farnesyltransferase (FTase) attaches a 15-carbon farnesyl moiety from farnesyl diphosphate while geranylgeranyltransferase type 1 (GGTase-I) links a 20-carbon geranylgeranyl group from geranylgeranyl diphosphate onto their substrates.^2^ These substrate proteins generally bear a canonical CaaX-box prenylation sequence where C is the prenylated cysteine and a and X can be any amino acid. However, recent studies have shown that proteins terminating in CXXXX or CXX may also potentially be substrates of prenylation.^3–5^ The third type of prenylation involves geranylgeranyltransferase type II (GGTase-II or RabGGTase) that usually appends two geranylgeranyl units on dual cysteine motifs on proteins, particularly those that belong to the Rab family. These substrates are typically recognized first by the rab escort protein 1 or 2 (REP-1/2) and modified by GGTase-II at their C-terminal cysteines in their -CCXX, -CXC, or -XXCC canonical sequences.^6^ Finally, the recently discovered GGTase-III appears to install a second isoprenoid geranylgeranyl moiety, onto to a pre-farnesylated protein substrate.^7,8^

Identifying prenylated proteins and the lipid-modified proteome in general has been an important research goal for more than a decade.^9,10^ Recent methods for high-throughput analysis of these lipid-modified proteins take advantage of chemical proteomics that allows for selective identification of protein substrates *via* an enrichment strategy.^11–13^ In this scheme, a bio-orthogonal probe (usually containing an alkyne moiety) that mimics the native form of a small molecule substrate used for enzymatic protein modification is added to cells and is incorporated by the host machinery into the target proteins. Alternatively, labeling of proteins can be achieved *in vitro* through the addition of the lipid probe and enzymes that catalyze the PTM.^14^ These labeled proteins present in lysates are then subjected to click reaction with a modified biotin for selective enrichment and subsequent proteomic identification of the modified proteins. Chemical proteomics approaches to identify prenylated proteins have been developed for more than a decade, wherein a bio-orthogonal isoprenoid analogue is synthesized and used to tag and identify the prenylated proteins in a given cell line.^10^ The goal of defining the prenylome has been a challenge, reflected by the lesser number of proteins identified in prenylomic studies compared to the total number of prenylation substrates derived from predictions and annotations. Recently, two independent studies reported 80 prenylated proteins identified in a given cell line, which consists of known and novel prenylation substrates.^11,12^ One study employed a dual labeling strategy where a farnesyl and a geranylgeranyl probe analogue were used, while the other used C15AlkOPP as a single probe that labels all classes of prenylation substrates. Despite the differences in these approaches, a comparable number of proteins were identified with slightly varying identities, particularly in the novel prenylated proteins discovered. In many cell lines that have been studied, one peculiar protein, ALDH9A1, has been consistently enriched. Here, we describe our discovery of ALDH9A1 as a potentially prenylated protein that has possible biological consequences in controlling this enzyme’s function. Through the use of tools in chemical and molecular biology, we characterized this potentially novel lipid modification and propose a mechanism via which this may occur in nature.

## Material and Methods

### General reagents and cell lines

COS-7 and HeLa cells were generously provided by Dr. Elizabeth Wattenberg at the University of Minnesota, USA. The FTI L-744,832 was purchased from Merck and GGTI-286 from CalBiochem. The RabGGTase inhibitors 1, 2, and 3 were prepared as previously described.^15,16^

### Plasmid preparation

A plasmid expressing human ALDH9A1 was purchased from Genecopoeia. The ALDH9A1 gene was inserted in a vector (EX-Z3075-M29) to contain an N-terminal GFP tag controlled by the CMV promoter. Plasmids were transformed in 100 µL of competent DH5α *E. coli* (New England Biolabs) following the manufacturer’s protocol. Transformed bacteria were plated in Luria broth (LB) agar plates containing ampicillin and colonies were isolated for bacterial cultures in 50 mL LB media. Cultures were grown overnight at 37 °C and bacteria were pelleted by centrifugation at 6000 *x g* for 20 mins at 4 °C. The bacterial pellets were suspended in sterile water and plasmids were extracted using a PureYield™ Plasmid Miniprep System (Promega) following the manufacturer’s protocol. Plasmid concentrations were determined by taking the absorbance at 260 and 280 nm using a NanoPhotometer P330 (IMPLEN) and using the equation: concentration (µg/mL) = (A_260_ – A_320_) x dilution factor x 50 µg/mL. Only good quality plasmids (A_260_/A_280_ ∼ 1.7 to 1.8) were used for subsequent transfections.

### Transfection, in-gel fluorescence and cellular imaging

COS-7 cells were grown in 100-mm culture plates under DMEM media (Gibco) supplemented with 10% fetal bovine serum (Gibco) and 1% penicillin-streptomycin (Gibco). Cells were passaged before reaching 80% confluency and repeated at least three times prior to transfection. A day before transfection, cells (1.6 × 10^6^) were seeded and grown overnight until 90% confluency was achieved. The media was removed and replaced with 10 mL of Opti-MEM™ Reduced Serum Media (Thermo Fisher Scientific). Plasmid DNA (15 µg) and 50 µL of Endofectin Max (Genecopoeia) were each diluted to 750 µL with Opti-MEM™ in separate tubes. The solutions were combined and added to the cells dropwise. After 8 hours post-transfection, 10 µM of C15AlkOPP, isoprenoid analogues, and inhibitors (when indicated) were added and allowed to incubate for 16 more hours. Cells were washed and collected by scraping and subjected to lysis in 1X PBS + 1% SDS as described in a previously reported protocol.^17^ Protein concentrations were determined using BCA assay (Thermo Scientific).

For in-gel fluorescence analysis, 100 µg of protein in lysate were used for the click reaction along with 25 µM TAMRA-N_3_, 1 mM TCEP, 0.1 mM TBTA, and 1 mM CuSO_4_. Labeled proteins were resolved in a 12% SDS-PAGE gel. TAMRA fluorescence (542/568 nm excitation/emission) was detected using a Typhoon FLA 9500 instrument (GE Healthcare). Gels were then stained with Coomassie blue and destained. For western blot analysis, proteins in gels were transferred to a PVDF membrane using a Mini Trans-Blot® Electrophoretic Transfer Cell (Bio-Rad) and blocked with 5% milk. Membranes were incubated with mouse anti-GFP at 1:5000 dilution in milk (ABclonal, #AE012) overnight followed by incubation with a secondary goat HRP-conjugated anti-mouse at 1:10000 dilution in milk (ABclonal, #AS003) and treated with Clarity™ ECL (Bio-Rad). Chemiluminescence was detected using iBright (Thermo Fisher). Gel images were processed using ImageJ.

For fluorescence cellular imaging, cells (500,000) were seeded in 35-mm glass-bottomed dish (Ibidi USA) and grown overnight until 90% confluency. The DMEM media was replaced with 2 mL of Opti-MEM™. DNA plasmids (2.5 ng in 125 µL Opti-MEM™) and Endofectin (7.5 µL diluted to 125 µL with Opti-MEM™) were mixed and added dropwise to the cells. After 24 hrs, the media was removed and cells were washed with cold 1X PBS (2 mL) twice, followed by staining with Hoechst 33342 (Thermo Scientific) and ER-Tracker™ Red (Invitrogen) following the manufacturers’ protocols. Cells were washed twice with PBS and visualized using a FluoView FV1000IX2 Inverted Confocal Microscope (Olympus) with a 60X objective. The .oib files were imported in Fiji and formatted.

### Site-directed mutagenesis

Human ALDH9A1 mutants were generated from the template plasmid (EX-Z3075-M29) described above using a QuickChange XL II Site-Directed Mutagenesis Kit (Agilent). Primers were designed with the corresponding mutations as prescribed by the QuickChange Primer Design tool (https://www.agilent.com/store/primerDesignProgram.jsp) with minor alterations at the 3’ end and the oligos were purchased from Integrated DNA Technologies. In brief, the template plasmid (50 ng) was incubated with the forward (125 ng) and reverse (125 ng) primers, dNTP mix, and PfuTurbo DNA Polymerase (2.5 U). PCR mutagenesis was carried out in an Arktik Thermal Cycler (Thermo Scientific) under the following thermal program: Initial denaturation (95 °C, 1 min); 18 cycles of denaturation (95 °C, 50 s), annealing (60 °C, 50 s), and extension (68 °C, 8 mins); final extension (68 °C, 7 mins). The resulting mixture was digested with DpnI and 5 µL of the mutagenesis reaction was transformed in XL10-Gold® Ultracompetent Cells (Agilent) and selected on amplicillin-containing LB-agar plates. Colonies were isolated, cultured and extracted with plasmids using PureYield™ Plasmid Miniprep System. Successful mutations were verified *via* Sanger-type DNA sequencing performed by the University of Minnesota Genomics Center. Purified plasmids were transformed in DH5α *E. coli*, cloned, purified, and assayed for concentration using NanoPhotometer P330 (IMPLEN).

### Mapping of the modification site

Expressed GFP-ALDH9A1 was enriched using anti-GFP beads following the manufacturer’s protocol for GFP-Trap (Chromotek). Briefly, COS-7 cells (∼1 × 10^6^) transfected with GFP-ALDH9A1 and treated with C15AlkOPP, FPP or GGPP were lysed in lysis buffer (100 uL, 10 mM Tris-HCl pH 7.5, 10 mM NaCl, 0.5 mM EDTA, 0.5% Nonidet™ P40 Substitute (Sigma-Aldrich)) by pipetting in and out in a 1-mL syringe with 16 gauge needle on ice. The soluble proteins were separated from the cellular debris via centrifugation (20,000 *x g* at 4 °C for 15 mins). Pre-washed GFP-Trap beads (25 uL) were added to the lysates and incubated for 1 h at 4 °C with head-to-tail mixing. The beads were then washed with wash buffer (10 mM Tris-HCl pH 7.5, 10 mM NaCl, 0.5 mM EDTA) and the remaining immobilized protein was reduced and digested in 50 mM Tris-HCl pH 7.5 containing 2 M urea, 10 ug/mL sequencing grade trypsin (Promega), 20 ug/ mL chymotrypsin (Promega), 1 mM CaCl_2_, and 0.5 mM TCEP for 30 min at 32 °C. The supernatant was collected using a spin trap (Pierce) and the beads were suspended in 50 mM Tris-HCl pH 67.5 containing 2 M urea and 5 mM iodoacetamide. The eluate was collected and pooled with the first eluate. Samples were dried via lyophilization and redissolved in 2% CH_3_CN in H_2_O with 0.1% HCO_2_H. The peptides were desalted up using STAGE tips following standard protocols. The recovered peptides were dried and redissolved in 0.1% HCO_2_H in H_2_O.

The samples were resolved using a reversed-phase column and analyzed with an Orbitrap Lumos instrument. Separation was performed in 5-30% of 0.1% HCO_2_H in CH_3_CN for 140 mins. A targeted database determined using Skyline was used corresponding to the *m/z* of the active site peptide resulting from tryptic and chymotryptic digestion with maximum of 3 missed cleavages. MS1 scans were collected at 60,000 resolution over 320-2000 *m/z* range with an AGC target of 500,000 and max IT of 50 ms. Dynamic exclusion was allowed for 90 s. HCD fragmentation was performed at NCE of 42% with 1.5 *m/z* isolation window, AGC of 50,000, and max IT of 200 ms. The .raw files were analyzed in pFind to search for the following modifications on cysteine residue: Carabamidomethylation (C_2_H_3_NO, 57.0215), C15Alk (C_18_H_26_O, 258.1984 Da), C15Alk-thioester (C_18_H_24_O_2_, 272.1776 Da), C15Alk-thiohemiacetal (C_18_H_26_O_2_, 274.1933 Da), farnesyl (C_15_H_24,_ 204.1878 Da), farnesyl ester (C_15_H_22_O, 218.1671 Da), farnesyl thiohemiacetal (C_15_H_24_O, 220.1827 Da), geranylgeranyl (C_20_H_32_, 272.2504 Da), geranylgeranyl ester (C_20_H_30_O, 286.2297 Da), and geranylgeranyl thiohemiacetal (C_20_H_32_O, 288.2453 Da). The neutral losses were also included. Other dynamic modifications include methionine oxidation (16.0000 Da) and arginine and glutamine deamidation (0.9840 Da). Matching spectra were exported and formatted for visualization.

## Results and Discussion

### Chemical proteomic analysis of prenylated proteins identified ALDH9A1

Recent advances in proteomic technologies have enabled high throughput profiling of pos-translationally modified (PTM) proteins of interest. In particular, chemical proteomic approaches involve an enrichment strategy to identify a set of modified proteins such as the prenylated proteome. Along with other groups, we have conducted prenylomic profiling across multiple cell lines and showed that the scope of prenylated proteins varies to some extent in these various cell types (Fig. 1).^12^ COS-7 (Fig. 1A) displayed the largest number of proteins profiled including novel prenylated proteins bearing the canonical CaaX-box motifs but were not previously identified in other prenylomic studies reported. We have found that the differential labeling of the prenylomes in these various cell lines may be attributed to the levels of expression of their prenyltransferases and cognate substrates or their ability for probe uptake.^12^

**Figure 1.**
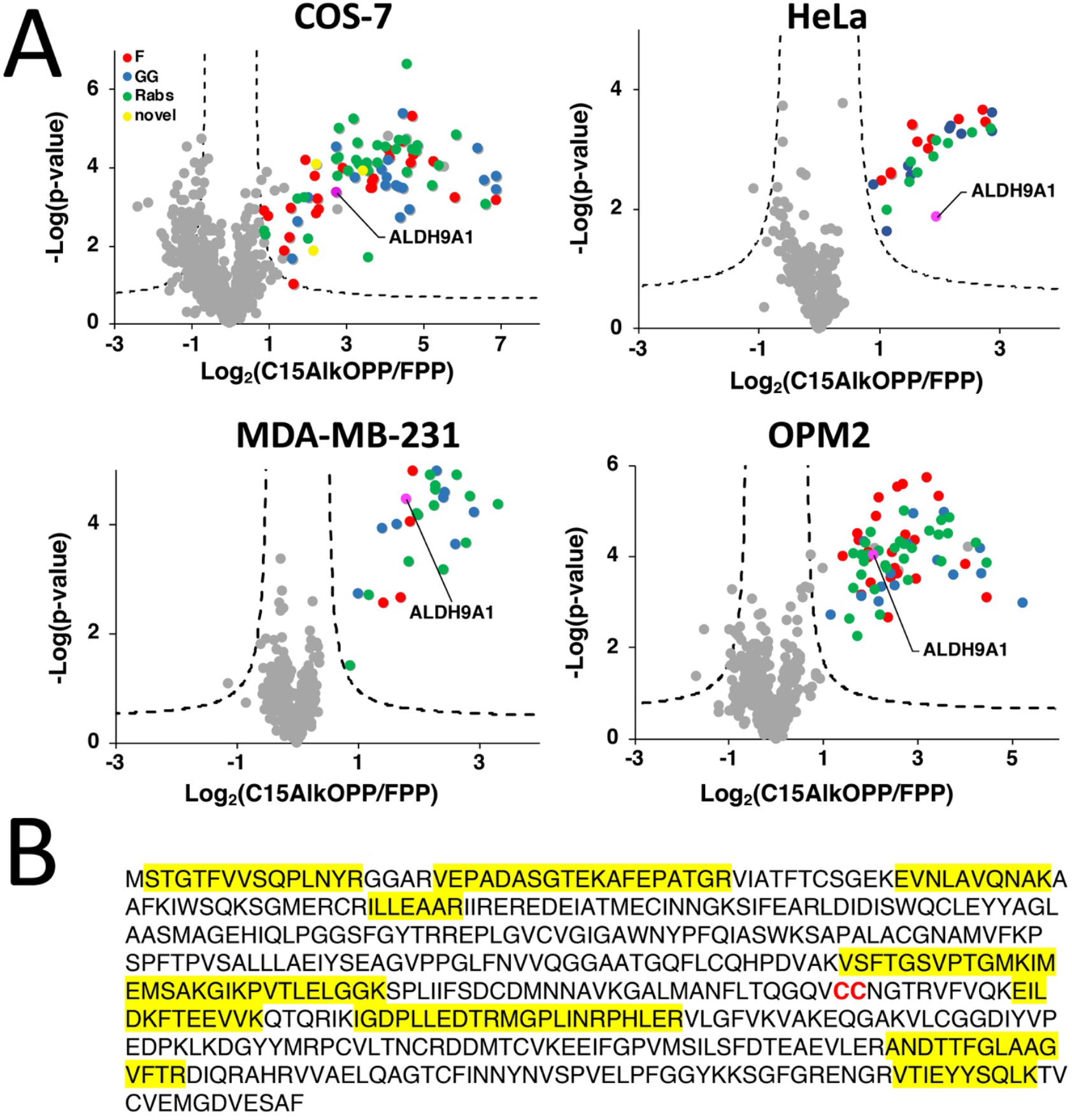
ALDH9A1 in consistently enriched in prenylomic analyses in various cell lines. **A.)** Volcano plots (FDR = 0.01, s0 = 0.5) of prenylomic analysis on various cell lines show thatALDH9A1 is consistently enriched. **B.)** The complete protein sequence of the human ALDH9A1. Highlighted in yellow are the tryptic peptides identified in the prenylomic analyses.

In our prenylomic studies, ALDH9A1, an aldehyde dehydrogenase, has been consistently enriched in a number of cell lines studied including COS-7, HeLa, MDA-MB-231, and OPM2 (Fig. 1A), as well as in studies from other groups such as in RAW264.7 cells.^12,18^ This enzyme does not bear a canonical prenylation motif but rather contains two adjacent cysteines (Cys288/Cys289) within its protein sequence, with Cys288 ascribed as a catalytic residue (Fig. 1B).^19,20^ We initially suspected that this protein may undergo proteolytic processing to reveal a cryptic C-terminal Cys-Cys motif (Cys288 and Cys289) that could serve as a potential substrate for dual prenylation by RabGGTase. However, examining the peptides identified in the proteomic analysis revealed that residues downstream of Cys288/Cys289 are enriched (Fig. 2B), suggesting that the intact full-length protein was isolated in the enrichment procedure. Therefore, labeling of ALDH9A1 with C15AlkOPP appears to deviate from the well-established rules for substrate recognition manifested by prenyltransferases.

**Figure 2.**
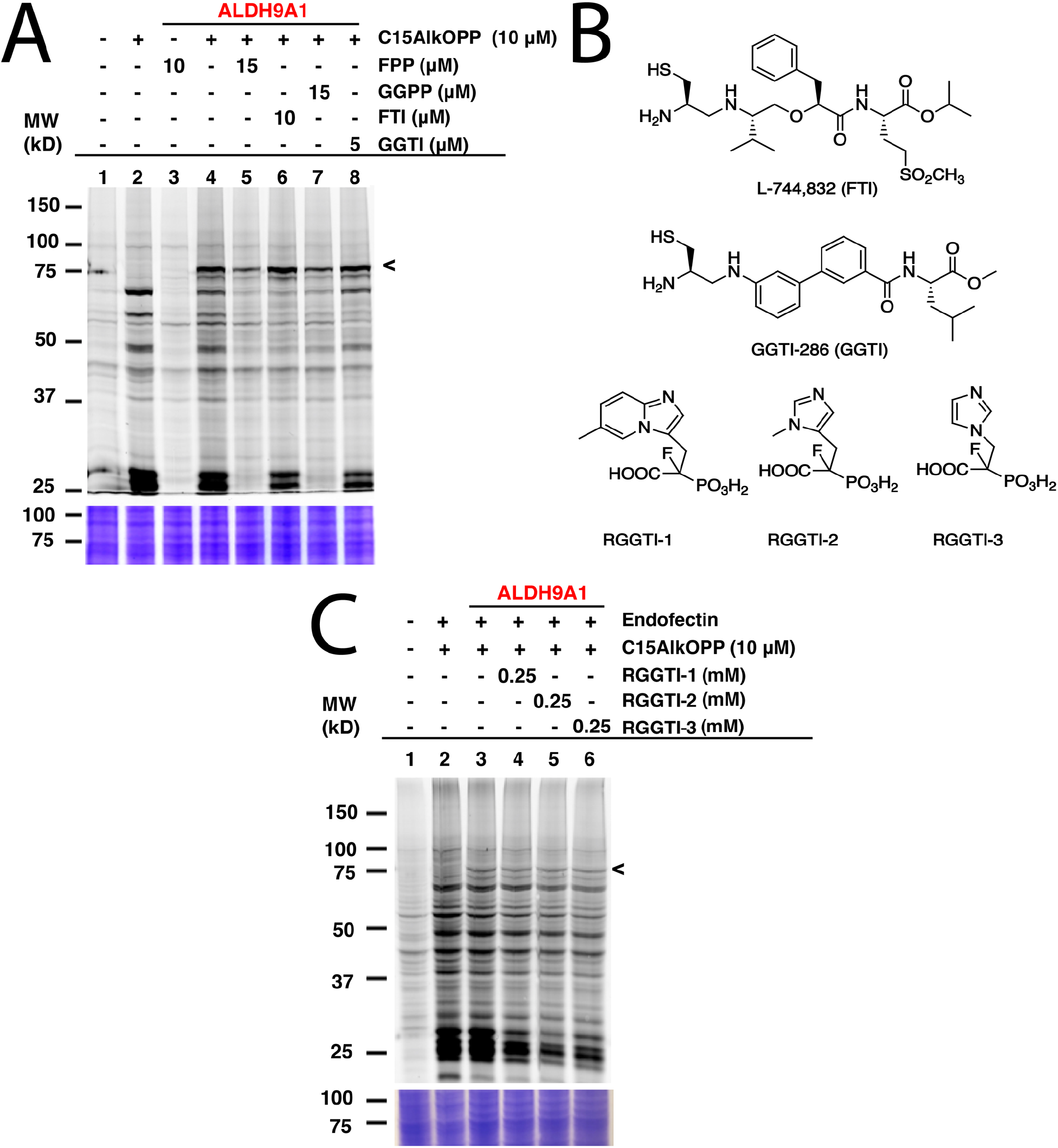
The C15AlkOPP probe labels ALDH9A1 in cells. **A.)** Labeling with isoprenoid probes with varying chain lengths in GFP-ALDH9A1 transfected in COS-7 cells. **B.)** Inhibition of the C15AlkOPP labeling with prenyltransferase inhibitors (FTI and GGTI) and competition with the native isoprenoids (FPP and GGPP). The C15AlkOPP labeling appears to be resistant in any of these inhibition/competition assays. **C.)** Structures of the prenyltransferase inhibitors used to inhibit the observed labeling on ALDH9A1. **D.)** Inhibition of the observed prenyl labeling on ALDH9A1 using RabGGTase inhibitors (RGGTIs). No apparent decrease in labeling of GFP-ALDH9A1 was observed although there was a significant decrease of labeling in the 25 kDa region where most Rab proteins migrate.

### C15AlkOPP labels ALDH9A1 but unresponsive to prenyltransferase inhibitors

In order to determine whether the C15AlkOPP probe indeed labels ALDH9A1 in the metabolic labeling experiments, the enzyme was overexpressed as a GFP fusion in COS-7 cells followed by subsequent metabolic labeling. A plasmid expressing ALDH9A1 with a C-terminal GFP tag (ALDH9A1-GFP) was initially transfected in COS-7 cells followed by incubation with C15AlkOPP post-transfection. By placing the GFP tag in a C-terminal position downstream of ALDH9A1, it might have been possible to evaluate whether endoproteolytic cleavage to expose a C-terminal prenylation motif was occurring. However, this approach did not result in successful expression of the protein (data not shown), perhaps owing to the fact that the C-terminal region of ALDH9A1 is important for homotetramerization that is essential for its stability.^19^ Instead, an N-terminal-tagged version (GFP-ALDH9A1) was used followed by metabolic labeling with C15AlkOPP and the lysates were subjected to click reaction with TAMRA-N3. Importantly, that construct displayed an intense fluorescent band near 75 kDa (Fig. 2A, lane 4), indicating successful labeling of the full-length intact protein. Since C15AlkOPP has the ability to label all three classes of prenylation substrates, it is not clear whether the ALDH9A1 is potentially farnesylated or geranylgeranylated.

To further establish the identity of the prenyl group modifying ALDH9A1, the C15AlkOPP probe was competed with the native isoprenoids. Co-treatment with 1.5-fold excess of FPP or GGPP with C15AlkOPP substantially but not completely abolished the labeling on ALDH9A1 (Fig. 2A, lanes 5 and 7). Under these conditions, virtually complete inhibition of the labeling of known prenylated proteins is readily achieved as indicated by the abolished signal in the 25 kDa region. Treatment with excess FPP inhibits both farnesylation and geranylgeranylation since FPP is elongated to GGPP by the GGPP synthase.^21^ As both FPP and GGPP competition reduces its labeling, ALDH9A1 may indeed be labeled with either isoprenoid. The prenyltransferases were also inhibited *in cellulo* by treating the transfected cells with known prenyltransferase inhibitors (Fig. 2B). Treatment with the FTase inhibitor L-744,832 (FTI, lane 6) or GGTase-I inhibitor GGTI-286 (GGTI, lane 8) did not significantly impact the labeling of ALDH9A1 under concentrations previously reported to induce observable inhibition of intracellular prenylation.^22,23^ In contrast, the reduction in the labeling of many of the normal prenylated proteins in the presence of FTI and GGTI is readily seen in lanes 6 and 8.

The absence of a CaaX-box motif in the full length ALDH9A1 suggests that it should not be a substrate of FTase and GGTase-I. However, this enzyme does contain two adjacent cysteines Cys288/Cys289 with Cys288 shown to be the catalytic residue for its aldehyde dehydrogenase function. As noted above, dual prenylation on adjacent cysteine residues is common among the Rab family of proteins that are substrates of RabGGTase, albeit only when they are near the C-terminus. Interestingly, a dual cysteine motif of Rab5b in *Plasmodium falciparum* was suggested to be prenylated although these putative prenylatable cysteines are 65 amino acids upstream from the C-terminus of that protein.^24^ Therefore the ability of known RabGGTase inhibitors (RRGTI-1-3) in diminishing the C15AlkOPP-labeling of GFP-ALDH9A1 was evaluated. RGGTI-1 and RGGTI-2 were previously shown to be effective in inhibiting Rab geranylgeranylation in HeLa cells (IC_50_ = 358 µM and 297 µM, respectively)^15,16^ while RGGTI-3 was less effective with IC_50_ = 850 µM. Treatment with these RGGT inhibitors in GFP-ALDH9A1-transfected COS-7 cells did not reduce the observed C15AlkOPP-labeling of GFP-ALDH9A1 (Fig. 2C). However, using this RGGTI concentration (250 µM), the prenylome labeling in the 25 kDa region was significantly diminished where most Rab prenylation substrates migrate (lanes 3 4, 5 and 6). If indeed RabGGTase prenylates ALDH9A1, a diminished labeling should have been observed in the presence of these RGGTIs. Therefore, ALDH9A1 does not appear to be a substrate of RabGGTase or any of the known prenyltransferases.

### Key residues in ALDH9A1 function are involved in isoprenoid labeling

The aldehyde dehydrogenase (ALDH) superfamily is a family of NAD^+^-dependent enzymes that catalyze the conversion of aldehydes to carboxylic acids with varying chain lengths and structures.^25–27^ ALDH9A1 in higher eukaryotes including humans is a homotetrameric enzyme that mediates the NAD^+^-dependent oxidation of many aldehydes such as betaine aldehyde, the carnitine precursor 4-trimethylaminobuteraldehyde (TMABAL), and the GABA precursor aminobutyraldehyde (ABAL).^28–30^ The general mechanism of catalysis for ALDH9A1 and for most of these ALDHs initially involves NAD^+^ co-factor binding that induces a conformational change resulting in activation of the thiol of a catalytic cysteine into a thiolate (Fig. 3A).^26^ In ALDH9A1, NAD^+^ binding in the coenzyme cavity is stabilized by many electrostatic interactions, which include the interaction between the pyrophosphate moiety of NAD^+^ that interacts with the residues Trp156, Ser233, and Thr236.^20^ The activated thiol then attacks the carbonyl of the aldehyde substrate, generating a thiohemiacetal intermediate stabilized by NH groups from the ALDH9A1 peptide main chain. A hydride transfer to NAD^+^ then follows, resulting in reduction of NAD^+^ to NADH and concomitant oxidation of the substrate to a thioester. This covalent intermediate then undergoes hydrolysis with the aid of a charged amino acid (usually Glu) in the active site to release the carboxylic acid product. The residue C288 is known to be the catalytic cysteine in ALDH9A1 while an adjacent Cys289 has no currently known function in this enzyme. A related yeast enzyme ALDH4 also contains this dual cysteine motif Cys324/Cys325 with Cys324 being the catalytic residue. A recent study has shown that Cys325 regulates the yeast enzyme’s function by forming a disulfide bond with Cys324, in order to promote an oxidative stress response.^31^ Similarly, the human ALDH1A1 exhibits the same disulfide bond formation between its active nucleophile Cys303 and the adjacent Cys302.^31^

**Figure 3.**
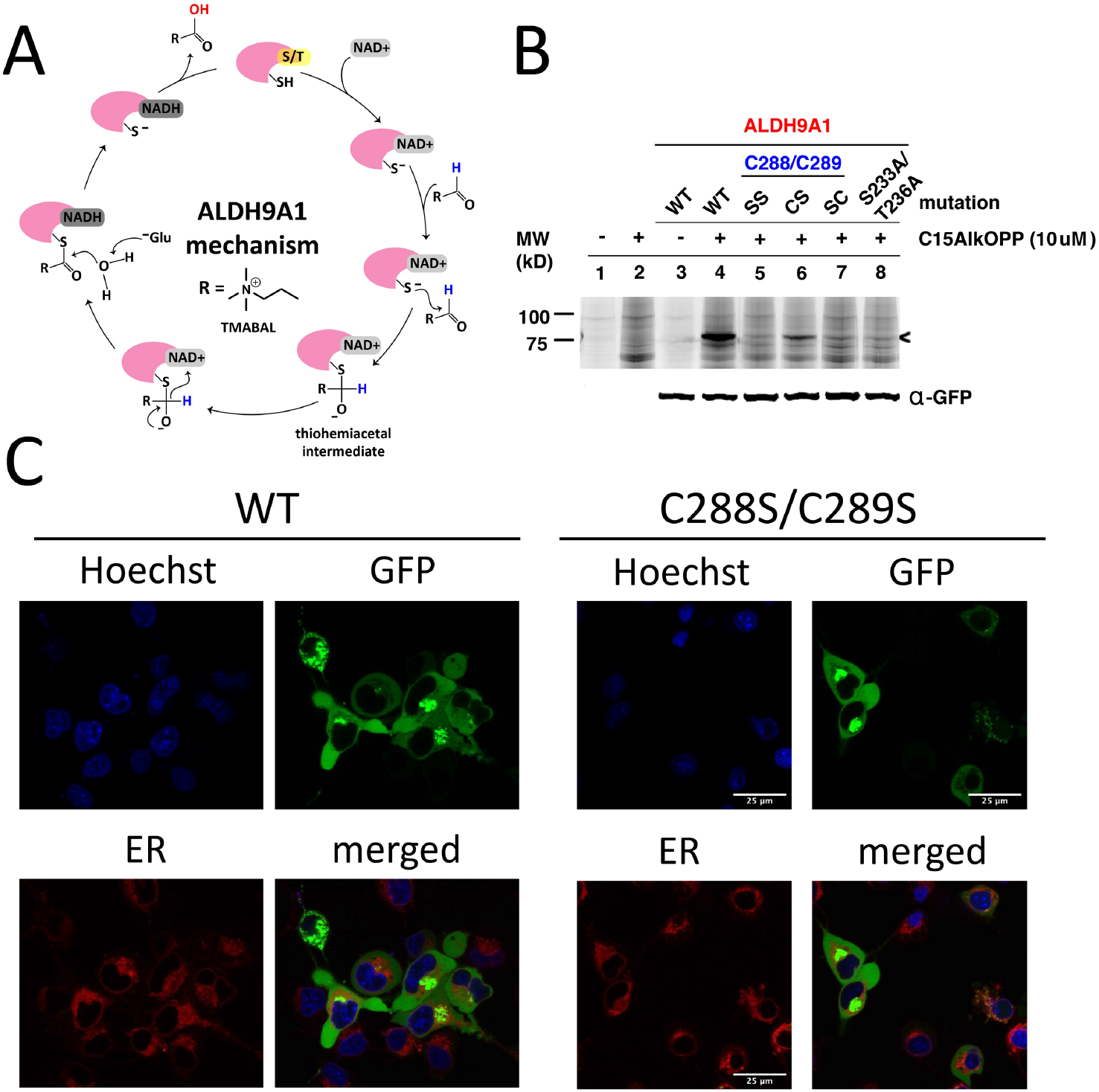
Essential amino acids in ALDH9A1 function influence prenyl labeling. **A**.) Mechanism of the ALDH9A1 activity. NAD^+^ binds to ALDH9A1 in the coenzyme cavity stabilized by many interacting residues including Ser233 and Thr236. The catalytic Cys288 is activated and attacks the carbonyl of the aldehyde substrate. A thiohemiacetal intermediate is formed and subsequently oxidized by NAD^+^ leaving a thioester bound substrate. A water molecule hydrolyzes the substrate assisted by an acidic amino acid (Glu). **B**.) In-gel fluorescence analysis of GFP-ALDH9A1-expressing COS-7 lysates metabolically labeled with C15AlkOPP. Labeling occurs on Cys288 and is affected by Cys289. Loss of the essential residues in the coenzyme cavity through double mutation Ser233A and Thr236A abolished labeling. **C**.) Fluorescent imaging of COS-7 cells transfected with wild-type or C288S/C289S mutant GFP-ALDH9A1. ALDH9A1 is distributed in the cytoplasm in both samples with observable puncta formation potentially resulted from aggregation. Hoechst, blue, nuclear stain; GFP, green, GFP-ALDH9A1; Endoplasmic reticulum (ER) tracker; red; ER.

Based on the mechanism described above, we decided to explore the role of some of the residues essential for ALDH9A1-mediated catalysis on labeling by C15AlkOPP. Therefore, site-directed mutagenesis was performed on the aforementioned protein construct to introduce mutations in some of these key residues and the resulting constructs transfected into COS-7 cells, followed by metabolic labeling with C15AlkOPP and click chemistry. Double mutation of the Cys288,Cys289 pair to serine residues (Cys288S,Cys289S, abbreviated SS) completely abolished labeling (Fig. 3B). Mutating the cysteine adjacent to the catalytic residue (C289S, CS) diminished the probe incorporation, whereas the mutating the catalytic cysteine only (C288S, SC) completely abrogated the signal. This suggests that probe labeling occurs on the catalytic Cys288 with Cys289 contributing in an indirect manner. As a control experiment, metabolic labeling was also performed in cells transfected with GFP-ALDH1A1 bearing this dual cysteine motif (Cys302/Cys303) but no labeling was observed (data not shown). We also examined the potential participation of NAD^+^ in the course of probe incorporation. Mutating residues Ser233 and Thr236, key amino acids essential for NAD^+^ binding, into alanine also completely abolished the observed labeling. Thus, these latter experiments suggest that NAD^+^ may be involved in this process of isoprenoid labeling of the catalytic Cys288 of ALDH9A1.

Protein prenylation is often associated with membrane targeting of the modified substrates, with the prenyl moiety serving as an anchor.^2^ Among the many protein prenylation substrates, mouse ALDH3B2 and ALDH3B3 are examples of aldehyde dehydrogenase enzymes known to be prenylated, both possessing the C-terminal sequence CTLL.^32^ Although both are prenylated, they differ in their cellular localization—ALDH3B3 localizes in the plasma membrane while ALDH3B2 resides in lipid droplets—influenced by some key residues in their corresponding C-termini.^32^ We therefore assessed the localization of the transfected GFP-ALDH9A1 fusion protein employed above and compared it with the corresponding double C288S/C289S mutant using fluorescence microscopy (Fig. 3C). There were no observable differences in the localization of the wild-type versus mutant. The protein appeared to be distributed across the cytoplasm, although discrete puncta were present in both samples. This protein species also did not localize in the endoplasmic reticulum (ER) where clusters of proteins reside during protein synthesis. Therefore, these puncta may be protein aggregates, which is a common phenomenon in protein overexpression in cells, particularly with proteins bearing a considerable number of cysteine residues (16 Cys in ALDH9A1).^33^ It is also possible that these puncta are lipid droplets, although the potential isoprenoid modification on ALDH9A1 may not necessarily influence this mechanism of localization. This hypothesis may be evaluated in future experiments.

### The isoprenoid modification is hydrolyzable

To gain insight into the origin of the isoprenoid incorporated into ALDH9A1, metabolic labeling experiments in COS-7 cells transfected with ALDH9A1 were performed with different isoprenoid analogues. The C15AlkOH probe (the alcohol form lacking the diphosphate group) produced appreciable labeling of ALDH9A1 although it was less compared to that observed with C15AlkOPP (Fig. 4A lane 3). It should be noted that isoprenyl alcohols readily become phosphorylated and incorporated into prenylated proteins.^34^ A previous report of using this analogue in profiling prenylated proteins identified ALDH9A1 as one of the highly enriched prenylated proteins.^18^ Thus, the isoprenoid modification on ALDH9A1 may be derived from the diphosphate analogue and that C15AlkOH may have been converted by host cell kinases to C15AlkOPP.^34^ We also evaluated the ability of the alkyne-modified analogue of farnesal (C15AlkCHO) and the methyl ester C15AlkCOOMe in labeling ALDH9A1 since the aldehyde and acid are both known isoprenoid metabolites in cells; of particular note, farnesal and geranylgeranial are known products of the deprenylating enzyme prenylcysteine oxidase (Pcyox 1).^35^ We rationalized that the neutral species C15AlkCOOMe could efficiently penetrate the plasma membrane and be hydrolyzed by cellular esterases to C15AlkCOOH, an analogue of FCOOH or GGCOOH. Neither of these analogues efficiently modified ALDH9A1 (Fig. 4A lanes 4 and 5) although some labeling was observed with C15AlkCHO. This reduced labeling was only present in wild-type ALDH9A1 and was abolished when mutations were introduced into the key residues (Fig. S1). It is possible that these analogues were bound to or reacted with serum proteins in the media and were not able to enter the cells since they both contain α,β-unsaturated carbonyl moieties that are known to readily react with nucleophiles.

**Figure 4.**
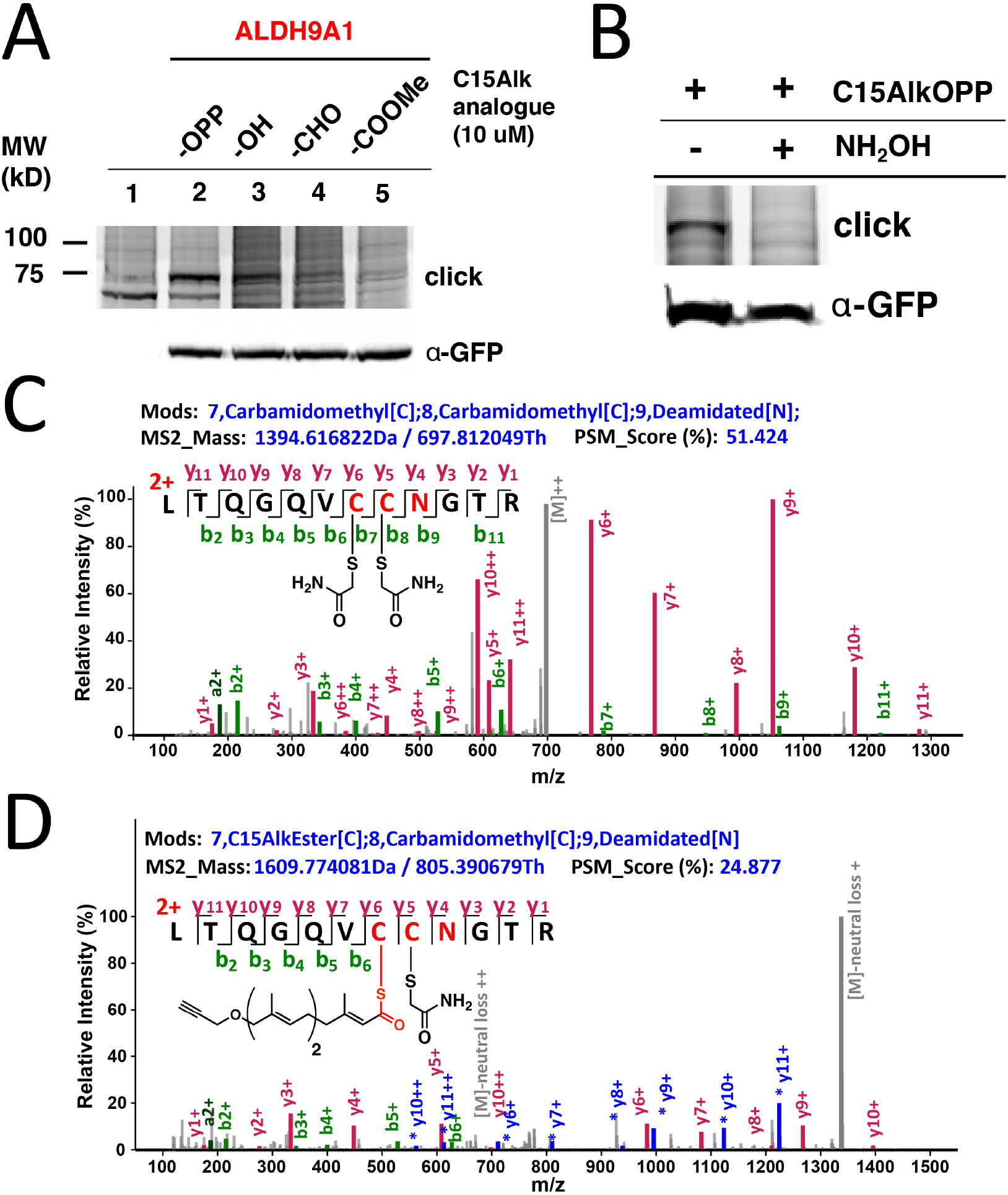
Hydrolyzable modification on ALDH9A1. **A**.) In-gel fluorescence analysis on lysates from COS-7 cells transfected with GFP-ALDH9A1 and treated with various alkyne-modified isoprenoid analogues. **B**.) Hydrolysis of the observed labeling on C15AlkOPP-labeled GFP-ALDh9A1 using NH_2_OH. **C**.) Mass spectrum of reduced/alkylated active site peptide of ALDH9A1. **D**.) Mass spectrum of the C15Alk-thioester-modified Cys288 of the active site peptide of ALDH9A1. Shown in blue and gray correspond to neutral loss of Alk-farnesoyl (C18H25O2).The colors of peaks are indicated for the parent ions (gray), y ions (red), y ions from parent peptide with the neutral loss (blue with *), and b ions (green).

While the aforementioned data with prenyltransferase inhibitors suggests that ALDH9A1 prenylation is not a process mediated directly by prenyltransferase, it does not rule out the conversion of C15AlkOPP into a variety of metabolites that could potentially result in subsequent ALDH9A1modification. Accordingly, we investigated whether this observed modification could be hydrolyzed by hydroxylamine. Basic hydrolysis with potent α-nucleophiles is commonly employed to distinguish reversible modifications such as *S-*palmitoylation on proteins since the thioester bond on palmitoylated cysteines is labile to hydroxylamine.^13^ Treatment of C15AlkOPP-labeled GFP-ALDH9A1-expressing COS-7 cells with hydroxylamine completely abolished the isoprenoid labeling (Fig. 4B). Thus, C15AlkOPP-mediated labeling appears to involve a hydrolyzable modification on ALDH9A1 instead of a thioether linkage that is resistant to hydrolysis that typically occurs in prenyltransferase-catalyzed protein modification.

The observed hydrolyzable modification on ALDH9A1 suggests that the isoprenoid modification may involve a thioesterified cysteine residue. Hence, we attempted to obtain a mass spectrum of the peptide modified with the alkyne-functionalized probe. COS-7 cells expressing the GFP-ALDH9A1 were treated with C15AlkOPP and the enzyme was immunoprecipitated using an anti-GFP resin. The immobilized protein was reduced, alkylated, and digested on-bead, generating tryptic peptides that were processed for LC-MS analysis. The reduced and alkylated form of the peptide containing Cys288 and Cys289 with deamidated N290 was efficiently detected with a retention time of 45.03 (Fig. 4C, Fig. S2A), indicating that some ALDH9A1 exists with free Cys288 and Cys289 thiols; however, some or all of this species could have arisen due to the hydrolytic instability of the thioester. Importantly, we also detected the C15Alk thioester-modified ALDH9A1 with deamidated N290 although that peptide gave less intense signals (Fig. 4D) and displayed a longer retention time (Fig. S2B), consistent with lipid-modified peptides. Deamidation is a commonly observed modification in MS analysis of peptides digested with trypsin.^36^ The detection of the fragment ions y6+, y7+, y9+ and y10+ (Table S1) are all consistent with the active site C288 thiol being modified with a thioester-linked isoprenoid. Numerous fragments originating from a neutral loss of the thioprenyl group were also observed (e.g. *y6+, *y7+, *y8+, *y9+, *y10+ and *y11+). Such neutral loss is commonly observed in the MS/MS fragmentation of thioacylated peptides such as those that are palmitoylated, as well as the thioether linkage in S-prenylated peptides.^37,38^ Efforts to repeat this experiment with FPP or GGPP were not successful. This may be related to the difficulty of detecting thioester-linked species in proteomic workflows due to their lability in the commonly used reduction and alkylation steps. With 16 cysteine residues, ALDH9A1 is particularly difficult to work with. Moreover, the presence of hydrophobic modifications from the isoprenoids on peptides commonly impacts their ionization efficiency in LC-MS analysis, thereby lowering the signal-to-noise ratio and complicating the detection of the lipid-modified peptide.^39^

Overall, a compelling case for the prenylation of ALDH9A1 is presented here. Initial work using chemical proteomics demonstrated that ALDH9A1 can be modified by the probe C15AlkOPP in at least 4 different cell types. Mutational analysis of the protein suggests that it is modified on Cys288 however experiments with various prenyltransferase inhibitors indicate that the modification is not catalyzed by a canonical prenyltransferase. The existence of a base-labile thioester and the requirement for an intact NAD^+^ binding site (inferred by mutagenesis) suggest a simple mechanism for this modification process where ALDH9A1 catalyzes the NAD^+^-dependent oxidation of a prenyl aldehyde to the corresponding thioester that is limited by a slow rate of hydrolysis resulting in accumulation of the covalently modified enzyme. Such a thioester species is a well-established intermediate in the mechanism catalyzed by aldehyde dehydrogenases^40^ and the slow hydrolysis of such acylated enzyme species is known to be the responsible for the ability of certain aldehydes to serve as covalent ALDH inhibitors.^41^ In this proposed mechanism, farnesal and/or geranylgeranial would originate from the prenylcysteine oxidase mediated deprenylation of farnesyl- or geranylgeranylcysteine. Previous work with ALDH9A1 demonstrated that the related isoprenoid, geranial is both an inhibitor and substrate that is slowly converted to geranic acid.^27,42^ That is directly analogous to what is proposed here with the exception that the isoprenoid is either five (farnesal) or ten carbons (geranylgeranial) longer. It is interesting to note that a substantial conformational change in the enzyme occurs upon nucleotide binding. In the NAD^+^ free form, the enzyme active site is open and the catalytic residue Cys288 is solvent exposed^19^ while in the presence of NAD+ that cysteine residue is sequestered (Fig. S3).^20^ In the later form, the hydrolysis of the putative thioester intermediate could be retarded, resulting in accumulation of the prenylated species.

An important question concerning these observations is do they have any biological significance? The ALDH superfamily constitutes an important class of enzymes that play key roles in modulating oxidative/electrophilic stress inside cells.^43^ In fact, dysregulation in their expression and function has been implicated in a variety of cancers and inhibitors specific to these enzymes have been developed for cancer therapeutics.^44^ Similar to sorbitol and alcohol dehydrogenases, ALDHs have been known to possess reactive nucleophilic cysteines in their active sites that drive the conversion of toxic aldehyde species to innocuous carboxylic acid derivatives, or in deactivating reactive aldehyde and oxygen species.^43,45^ The recent developments in chemical proteomic strategies have allowed for profiling of proteins bearing reactive cysteines in which several ALDHs have been identified.^46–48^ In particular, ALDH9A1 has been consistently identified in these chemical proteomic studies, underscoring the high reactivity of its active Cys288 residue. Another salient characteristic of ALDH9A1 is the presence Cys289 adjacent to the nucleophilic Cys288. Other ALDHs also contain this dual (even triple) cysteine motif: ALDH2 (Cys318/Cys319*/Cys320), ALDH1A1 (Cys302/Cys303*), ALDH1A2 (Cys319/Cys320*), ALDH1A3 (Cys313/Cys314*), and ALDH1B1 (Cys318/Cys319*/Cys320), where * denotes the catalytic cysteine. Among these ALDHs, only ALDH9A1 was consistently enriched in our prenylomic analyses. Previous studies in ALDH1A1 have shown that Cys302 adjacent to the nucleophilic Cys303 may control its activity through disulfide bond formation and is potentially important for cellular response to oxidative stress.^31^ Prenylation of Cys288 in ALDH9A1 may be a related mechanism for controlling the activity of that enzyme. Interestingly, we recently identified ALDH9A1 as a protein with increased levels of prenylation in Alzheimer’s disease.^49^ Since prenylation of ALDH9A1 on the catalytic residue Cys288 must inhibit its activity, this observation may provide a link between oxidative stress, neuroinflammation and Alzheimer’s disease.^50^

## Conclusion

In this work, we described the potential isoprenoid modification on ALDH9A1 discovered through chemical proteomics. Through the use of tools from chemical and molecular biology, we demonstrated that this isoprenoid labeling on ALDH9A1 is robust but reversible. This potentially new type of isoprenoid modification opens new doors into the chemistry of lipid modifications on proteins, particularly in modifying the catalytic residue of an enzyme that may have functional roles in regulating its activity. The experiments described here set the stage for future work aimed at understanding this unusual post-translational modification. More studies of this novel protein modification are therefore anticipated.

## Data Availability

The raw mass spectrometry files for the proteomics data have been deposited to the ProteomeXchange Consortium via the PRIDE partner repository with the dataset identifier PXD036365.

## Acknowledgements

The authors thank Dr. Katarzyna Blazewska for generously providing the RabGGTase inhibitors used in this study. The authors acknowledge the Minnesota Supercomputing Institute (MSI) at the University of Minnesota for providing resources that contributed to the prenylomic research results reported within this paper (http://www.msi.umn.edu) and Dr. Yingchun Zhao and Dr. Peter Villalta for the assistance with proteomic data collection in the Analytical Biochemistry Shared Resource of the Masonic Cancer Center, designated by the National Cancer Institute and supported by P30 CA077598

## Author information

### Contributions

KF performed prenylomic analysis, cellular imaging, transfection and mutagenesis experiments. GS and SA assisted with some gel experiments and enzyme assays.

KS, LL and MD conceptualized, advised, and guided project development. KS and MD wrote the paper with input and guidance from all authors.

KS and MD interpreted the data.

All authors critically read and reviewed manuscript.

