## Supporting Information for "Thinking outside the CaaX-box: an unusual reversible prenylation on ALDH9A1"

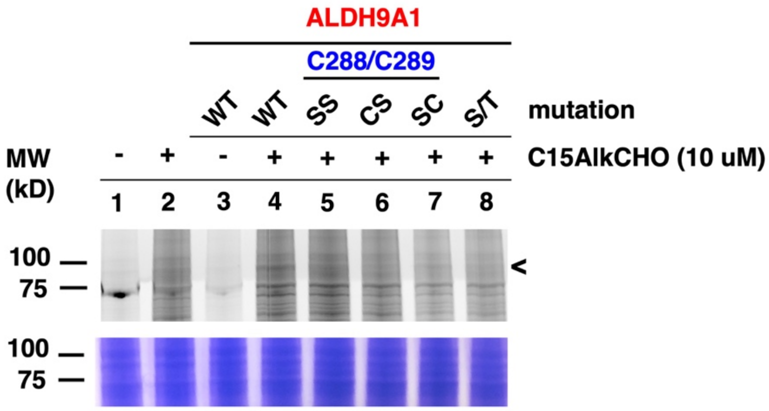

**Figure S1.** In-gel fluorescence analysis of GFP-ALDH9A1-expressing COS-7 lysates metabolically labeled with C15AlkCHO.

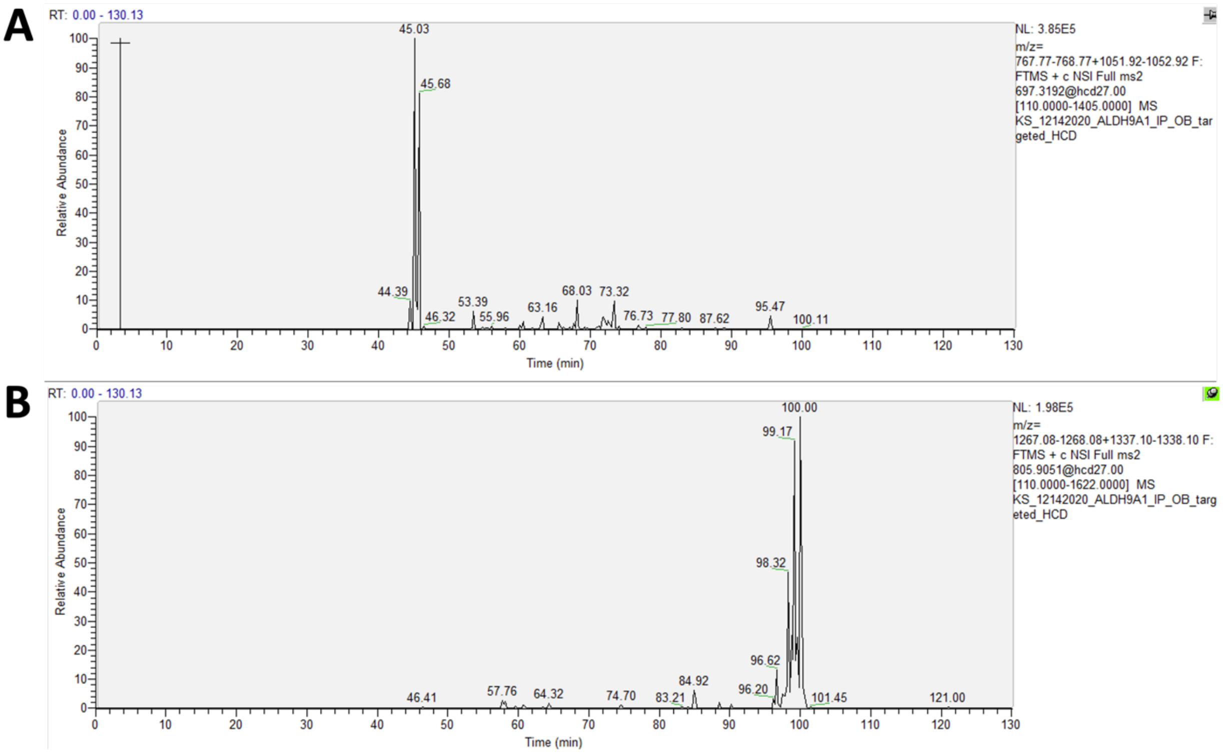

**Figure S2**. Extracted ion chromatograms corresponding the unmodified (A) and C15Alkthioester-modified (B) active site peptide of ALDH9A1.

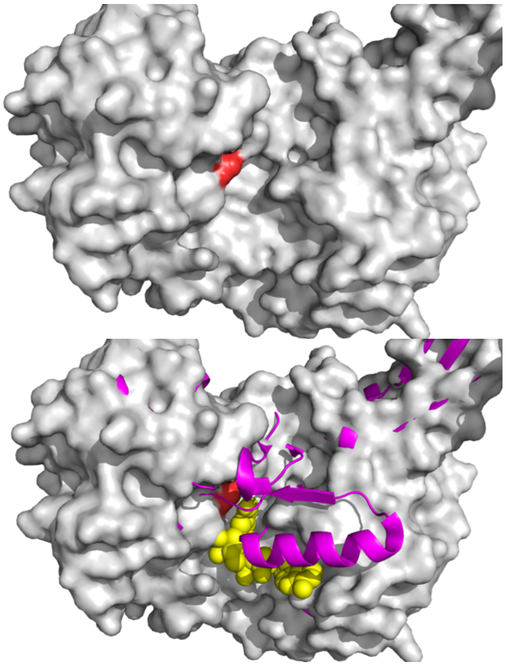

**Figure S3.** Structures of ALDH9A1**.** Top: Structure of enzyme in the absence of NAD+(pdb code 6qap). Bottom: Structure of enzyme in the presence of NAD+ (pdb code 6vr6). Color scheme: enzyme in the absence of NAD+ (grey); Cys288 (red); Enzyme in the presence of NAD+(magenta); NAD+(yellow).

**Table S1.** Fragment ions of the ALDH9A1 active site tryptic peptide obtained without and with thioester modification.

| Ion Type | C288(Cam),C289(Cam)[a] | | | C288(C15Alkthioester),C289(Cam)[a] | | |
|---|---|---|---|---|---|---|
| | m/z | intensity | m/z error | m/z | intensity | m/z error |
| b2+ | 215.1390 | 14.6832 | -1.1526 | 215.1390 | 4.7479 | -0.0835 |
| b3+ | 343.1976 | 5.7959 | 0.3224 | 343.1976 | 1.7322 | 2.0123 |
| b4+ | 400.2190 | 6.2064 | -0.2760 | 400.2190 | 2.1554 | 2.0227 |
| b5+ | 528.2776 | 10.2154 | -0.4011 | 528.2776 | 3.6567 | 1.1133 |
| b6+ | 627.3460 | 10.8375 | -0.6066 | 627.3460 | 3.3977 | 2.1191 |
| b7+ | 787.3767 | 1.9539 | -0.1713 | 1002.5328 | - | - |
| b8+ | 947.4073 | 1.0288 | 2.2280 | 1162.5635 | - | - |
| b9+ | 1062.4342 | 3.9907 | -1.8800 | 1277.5904 | - | - |
| b11+ | 1220.5034 | 1.1269 | 0.3260 | 1435.6595 | - | - |
| b11++ | 610.7553 | 0.5195 | -16.2246 | 718.3334 | - | - |
| y1+ | 175.1189 | 5.0454 | -1.5034 | 175.1189 | 1.9810 | -0.3613 |
| y2+ | 276.1666 | 2.2817 | 0.1562 | 276.1666 | 1.6655 | 1.3511 |
| y3+ | 333.1881 | 18.8114 | 0.0061 | 333.1881 | 15.4811 | 0.9365 |
| y4+ | 448.2150 | 8.3148 | 0.0978 | 448.2150 | 10.3656 | 0.7002 |
| y5+ | 608.2457 | 23.3810 | -0.4940 | 608.2457 | 11.1590 | 0.6075 |
| y6+ | 768.2763 | 91.2758 | -0.7091 | 983.4325 | 11.1932 | 0.1629 |
| y6++ | 384.6418 | 1.8886 | -0.8076 | 492.2199 | - | - |
| y7+ | 867.3447 | 60.3675 | -0.6842 | 1082.5009 | 7.6610 | -0.9131 |
| y7++ | 434.1760 | 1.1442 | -0.4260 | 541.7541 | - | - |
| y8+ | 995.4033 | 22.1949 | -0.6378 | 1210.5594 | 1.8784 | 0.0084 |
| y8++ | 498.2053 | 1.8408 | 0.0187 | 605.7834 | 0.0000 | 0.0000 |
| y9+ | 1052.4247 | 100.0000 | -1.4500 | 1267.5809 | 10.4304 | 0.8907 |
| y9++ | 526.7160 | 1.8801 | -0.4770 | 634.2941 | - | - |
| y10+ | 1180.4833 | 28.8661 | -1.8953 | 1395.6395 | 1.7641 | -0.9260 |
| y10++ | 590.7453 | 66.1492 | -0.2995 | 698.3234 | 0.7228 | -3.6866 |
| y11+ | 1281.5310 | 2.6356 | -3.6058 | 1496.6871 | - | - |
| y11++ | 641.2691 | 32.1587 | 0.1501 | 748.8472 | - | - |

[a]C288(Cam)C289(Cam) refers to the peptide where both cysteine residues have been alkylated with carboxamidomethyl groups at positions 288 and 289, respective whereas C288(C15Alkthioester)C289(Cam) refers to the peptide where Cys288 has a thioester and Cys289 has a carboxamidomethyl group.

**Sequences of primers used for mutation**

C288S/C289S:

Forward: 5'-AAT  ACT CTT GTG CCA TTA GAG GAA ACC TGG CCT TGT GTG A-3'
Reverse: 5'-TCA CAC AAG GCC AGG TTT CCT CTA ATG GCA CAA GAG TAT T-3'

CS: C289S

Forward: 5'-ACA CAA GGC CAG GTT TGC TCT AAT GGC ACA AGA GTA TTG-3'
Reverse: 5'-AAT ACT CTT GTG CCA TTA GAG CAA ACC TGG CCT TGT GTG AG-3'

SC: C288S

Forward: 5'-CTC ACA CAA GGC CAG GTT AGC TGT AAT GGC ACA AG-3'
Reverse: 5'-CTT GTG CCA TTA CAG CTA ACC TGG CCT TGT GTG AG-3'
